## Supplementary material for "Redox Regulation of Brain Selective Kinases BRSK1/2: Implications for Dynamic Control of the Eukaryotic AMPK family through Cys-based mechanisms"

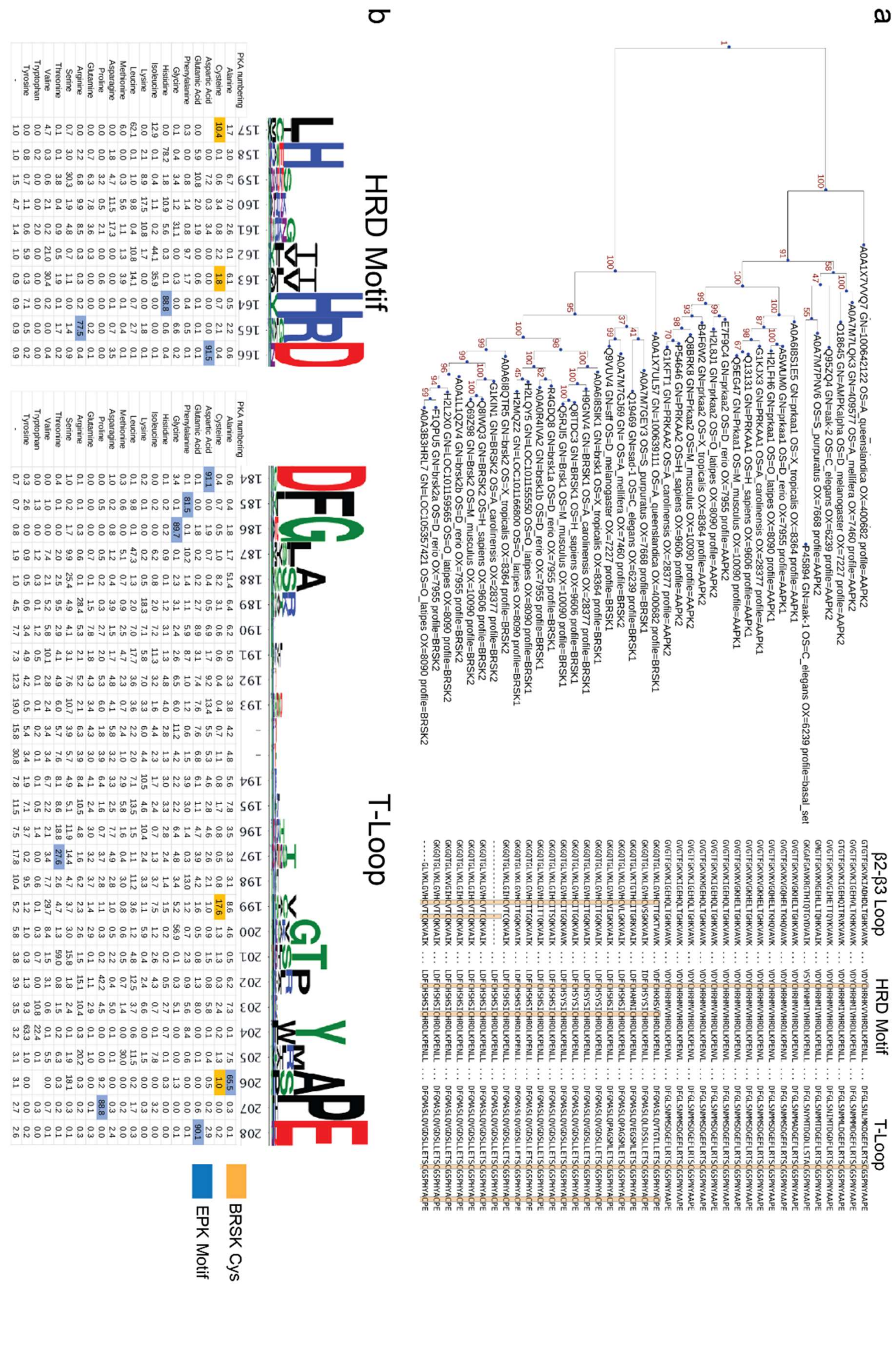
a
b
β2-β3 Loop
HRD Motif
T-Loop
BRSK Cys
EPK Motif

Supplementary Figure 1: Phylogenetic analysis of BRSK and the ARK family. (a) Phylogenetic analysis of the ARK family reveals that the closest relative of BRSK kinases is AMPK. The number of cysteines in the kinase domain of BRSKs increases relative to AMPK. (b) Sequence alignment and relative amino acid composition of the activation segment of ePKs (top). Data is presented as HMM (hidden Markov models) Sequence Logos. Table (bottom) depicts the frequency of an amino acid at each position along the Catalytic and T-Loop. Key Cys residues are highlighted in orange; residues highly conserved in ePK canonical kinase motifs are highlighted in blue.

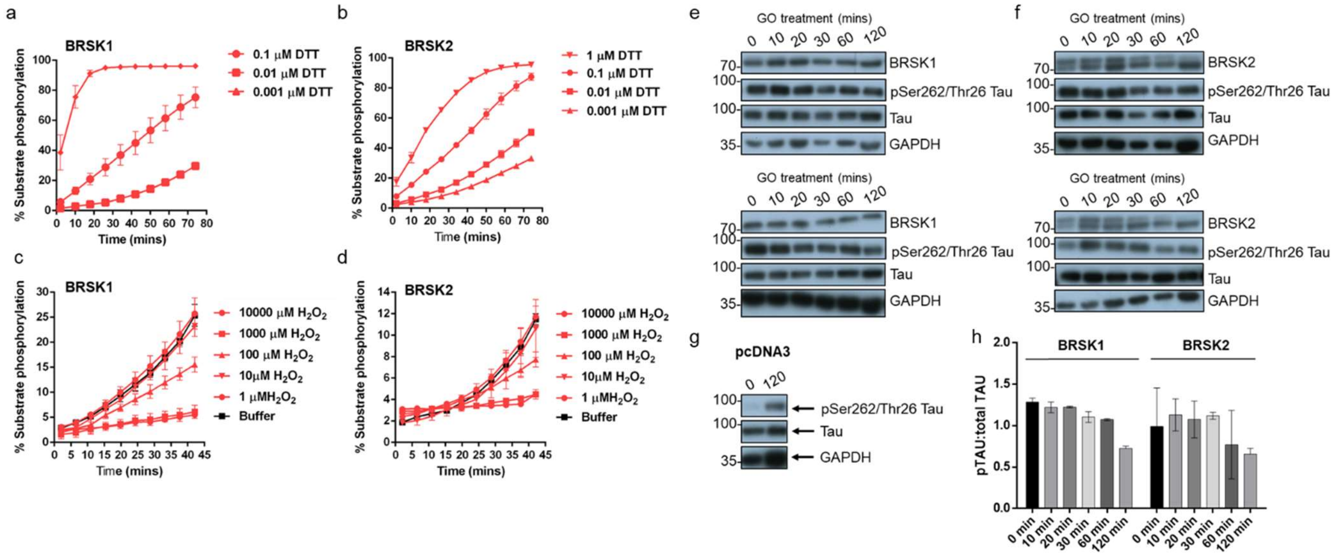


Supplementary Figure 2: Redox regulation of BRSK1 & 2. (a-d) Real-time phosphorylation of fluorescent AMARA peptide by full length BRSK1 and 2 (200 ng). BRSK proteins were incubated with buffer or the indicated concentrations of DTT or $H_2O_2$. Rates of BRSK activity were calculated as pmol per min phosphate incorporation and are presented in Fig 1. Data shown here is a subset of the conditions shown in Fig 1 (mean and SD from three repeats). (e-h) Time dependent loss of pTAU by incubation of HEK-293T cells with 2U/ml glucose oxidase (GO). Cells were transiently co-transfected with EGFP-Tau and either (e) BRSK1, (f) BRSK2 or (g) empty vector (pcDNA3). Data shown is WB analysis from 2 independent repeats. (h) pTau:Tau signals calculated with ImageJ. Data shown is mean and SD, calculated from (e) and (f).

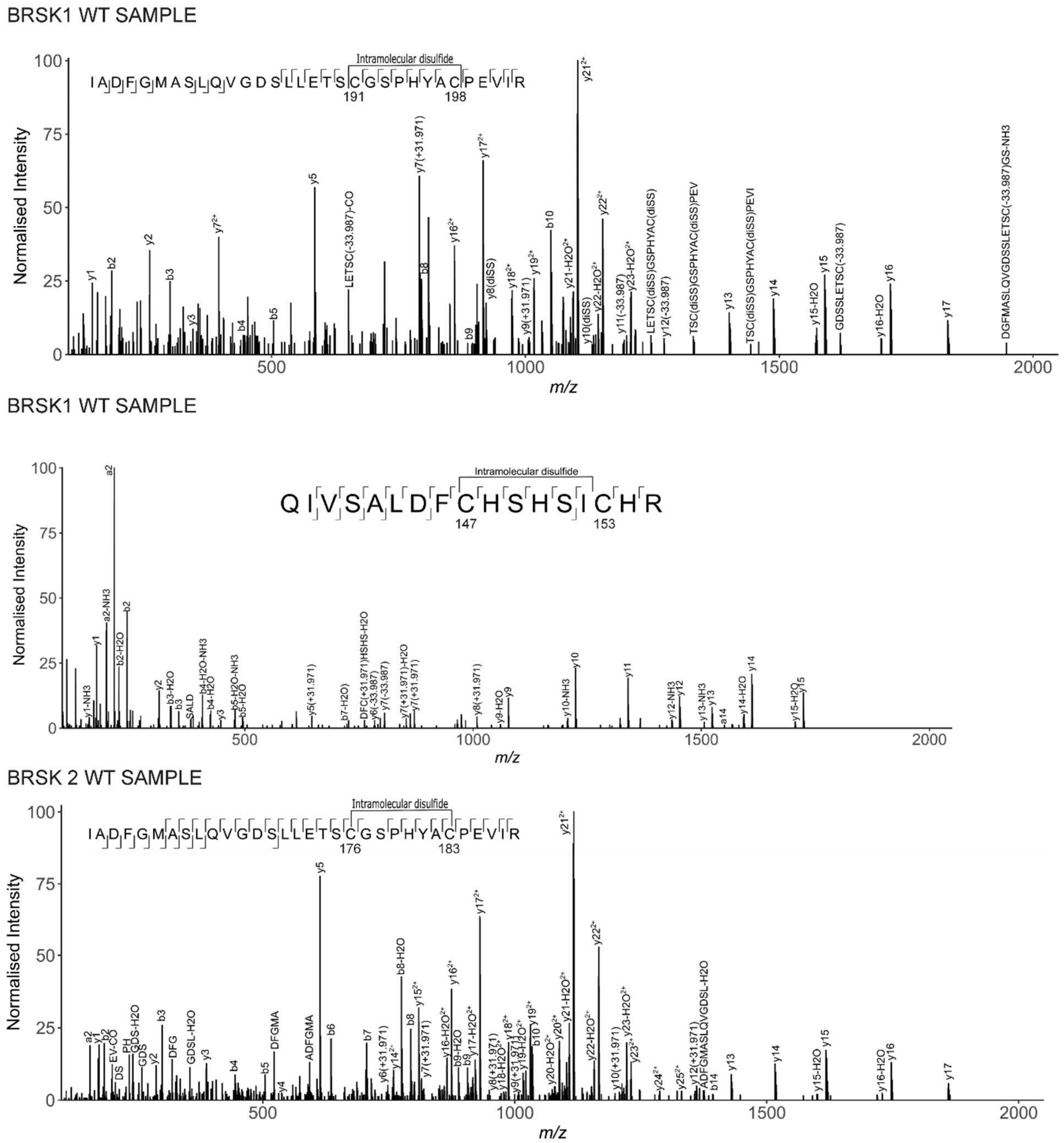


Supplementary Figure 3: LC-MS/MS Analysis of BRSK1/2 catalytic domains. LC-MS/MS reveals intramolecular disulfide bonds in the kinase domains of BRSK1 and 2 purified from *E. coli*.

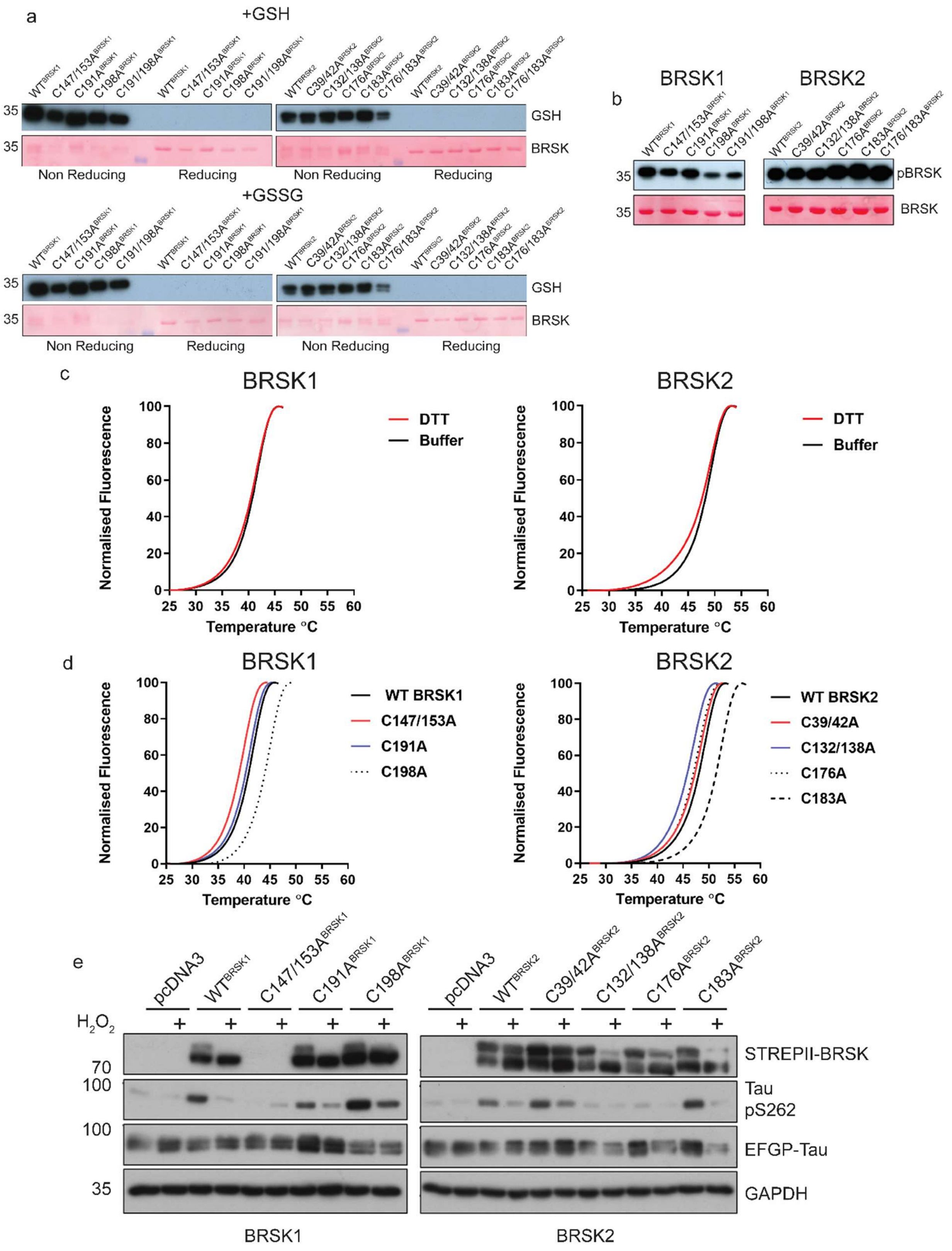
a
+GSH
WT BRSK1
C147/153A BRSK1
C191A BRSK1
C198A BRSK1
C191/198A BRSK1
WT BRSK2
C39/42A BRSK2
C132/138A BRSK2
C176A BRSK2
C183A BRSK2
C176/183A BRSK2
35
GSH
BRSK
Non Reducing
Reducing
+GSSG
b
BRSK1
BRSK2
pBRSK
c
BRSK1
BRSK2
DTT
Buffer
Normalised Fluorescence
Temperature °C
d
WT BRSK1
C147/153A
C191A
C198A
WT BRSK2
C39/42A
C132/138A
C176A
C183A
e
pcDNA3
$H_2O_2$
+
70
100
STREPII-BRSK
Tau pS262
EFGP-Tau
GAPDH

Supplementary Figure 4: Biochemical analysis of BRSK Cys-to Ala mutants. (a) Immunoblot of in vitro glutathionylation of BRSK kinase domains. (b) Immunoblot showing LKB1-dependent phosphorylation of BRSK kinase domain proteins. (c) Thermal denaturation curves of BRSK catalytic domain proteins in the presence or absence of 10 mM DTT. (d) Thermal denaturation curves of BRSK catalytic domain cysteine to alanine mutants. (e) Representative immunoblot of EGFP-Tau co-expressed with full length, WT and Cys-to-Ala mutants of BRSK1 and BRSK2. Transiently transfected HEK-293T cells were treated with or without 10 mM $H_2O_2$ for 10 mins.

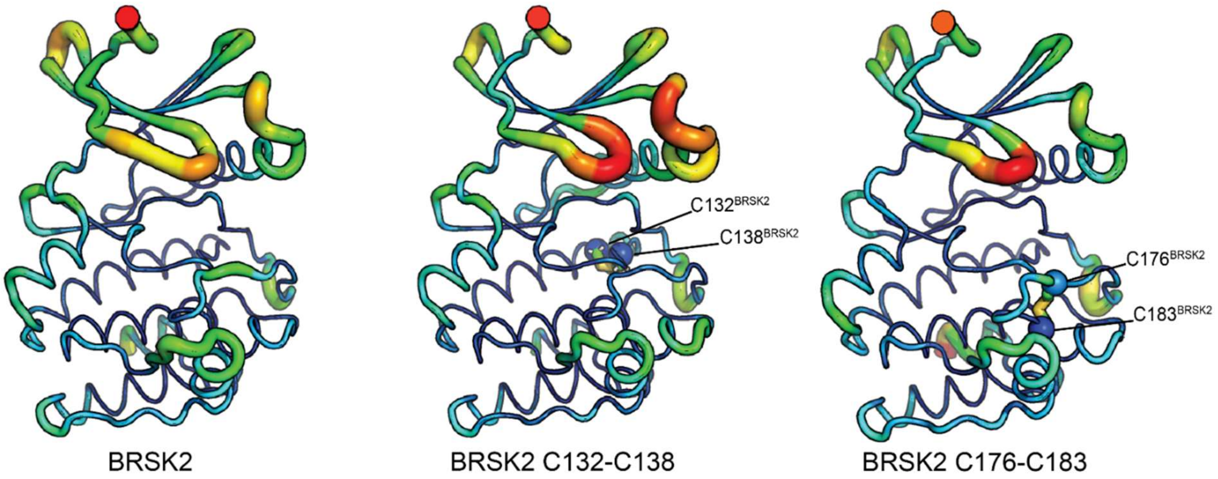


Supplementary Figure 5: Molecular Dynamics Simulations of intramolecular disulfide bonds. Simulations incorporating disulfide bonds identified in MS/MS experiments. RMSF was calculated based on three 100 ns GROMACS molecular dynamics simulations. Higher mobility is indicated by warmer colors and thickness of representation.

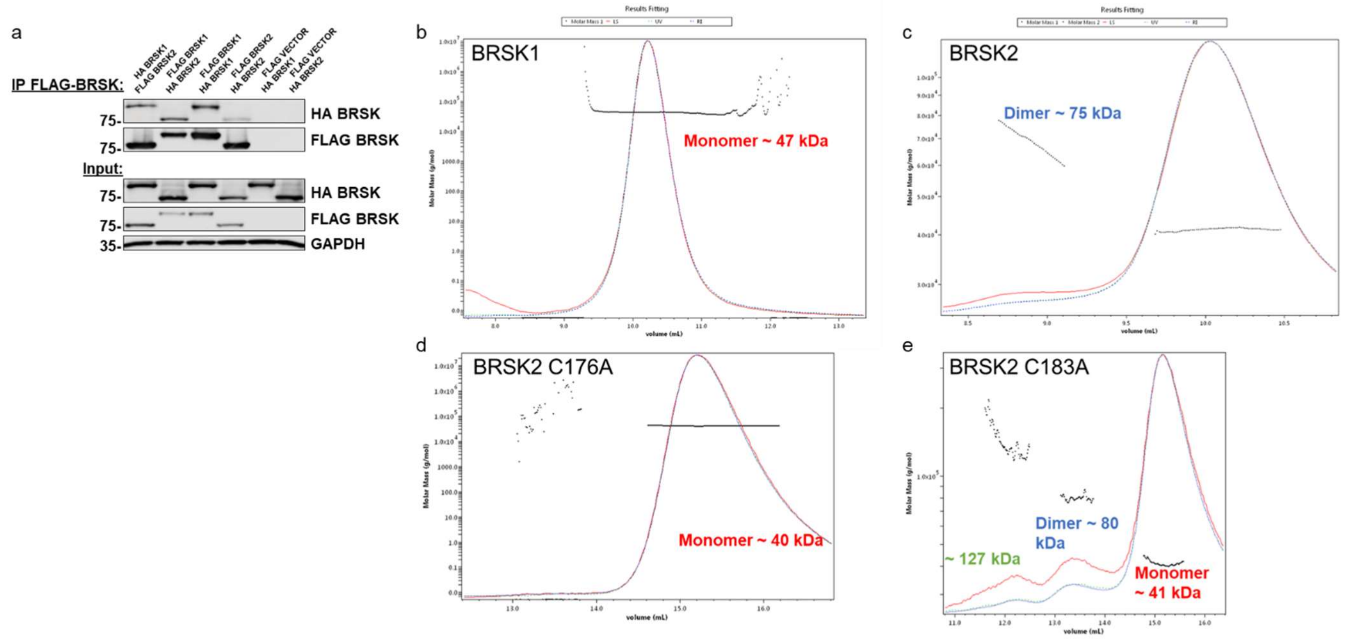


Supplementary Figure 6: Evidence for limited BRKS dimer species. (a) Co-immunoprecipitation of HA-BRSK1/2 with immunoprecipitated FLAG-BRSK1/2 expressed in HEK-293T cells. SEC-MALS analysis of WT (b) BRSK1 and (c) 2, and (d) C176A and (e) C183A BRSK2 kinase domains in solution, performed in the absence of reducing agents.
